# Single-cell RNA-seq Reveals Early Transcriptional Programs of the Maternal to Zygote Transition in Mice and Rats

**DOI:** 10.1101/2024.09.30.615942

**Authors:** Yangqi Su, Jane Kouranova, Jonathan Shea, Xiaoxia Cui, Zhengchang Su

## Abstract

The maternal to zygote transition in mammals has been an area of intensive research over the past few decades, with an ever-changing landscape of understanding that has accompanied the rapid development of cell-profiling technology. Utilizing a full-length single cell RNA-seq protocol, we profiled mature oocytes and zygotes of mice and rats to uncover elusive transcriptomic dynamics during the maternal to zygote transition. We note the existence of early gene expression of crucial zygotic development pathways in the mouse zygote while revealing a similar chain of events occurring in the rat zygote. We further observe an increase in nascent and intergenic transcription in both species. Moreover, we find subtle but pervasive signals of differential transcript usage in genes related to epigenetic regulation occurring in both species. In terms of post-transcriptional modifications, we find distinct profiles of alternative polyadenylation between zygotes and oocytes in both species, particularly, in genes related to cell cycle processes within the zygotes of mice. Finally, although a more dynamic transcriptomic landscape exists in the mouse zygote, the rat zygote also displays similar transcriptomic features, suggesting that minor zygotic activation in rat may occur earlier than originally thought.

## 1. Introduction

Fusion of the oocyte and sperm forming the zygote marks the inception of all mammalian embryogenesis, during which the male and female genome come together to jumpstart the developmental process^1^. Despite providing half the DNA and being crucial for oocyte activation, the sperm brings few components in the zygote and thus provides smaller resources in the early stages of embryogenesis than the oocyte. It is therefore mainly the contents of the maternal gamete that set up the suitable environment for successful polyspermy prevention and zygote genome activation (ZGA). During ZGA the control of cellular processes is gradually let go mainly from the products of the maternal genome and transferred to products of the zygote genome^2,3^. This process, also known as maternal to zygote transition (MZT) ^4^, has long been the subject of interest for researchers. Most foremost, the oocyte is a highly differentiated cell that quickly acquires pluripotency after fertilization. Though much research has been conducted in this area, it is still an elusive topic, and many intricacies remain to be uncovered.

An aspect that has been observed in MZT of all mammals has been the silence of the transcriptome in mature oocytes prior to fertilization up until ZGA after fertilization, which initiates the events of embryogenesis^5^. The length of this period of transcriptional silence varies in different species. This quiescent period that is conserved among species has not been fully understood, particularly, the mechanisms underlying ZGA may be different for each species^6^. Nonetheless, in all the mammalian species the machineries that carry on the progression of meiosis and remodel the genome depend solely on the maternal mRNAs accumulated during oocyte development^5,6^. Thus prior to ZGA, there is also translation and degradation of many maternal mRNAs^7,8^.

The mouse (*Mus musculus*) and rat (*Rattus norvegicus*) are two of the most used model mammals in biomedical research. Many aspects of the two organisms are very similar, such as genome size and number of protein-coding genes. It is estimated that at least 90% of genes in the mouse genome have at least a homolog in the rat genome, and vice versa ^9^. However, there remain considerable differences in the biology of the two organisms including reproduction^10^. For instance, it is known that ZGA occurs in 2 waves in mice with a ‘minor ZGA’ in the late 1-cell stage and a ‘major ZGA’ in 2-cell stage. In contrast, it was reported that the onset of embryonic transcription activity in rats occur in late 2-cell stage^11,12^. However, even though major ZGA is known to occur at the 8-cell stage in human embryos, recent research has suggested that minor ZGAs occur in 1-cell human zygotes as well, though to a lesser degree than that of mice^13^. Thus, it is possible that similar transcriptional events might also occur in 1-cell rat zygotes.

Research into transcriptional events occurring during minor ZGA in mice has revealed insightful results. Early studies in mice using brUTP incorporation found that transcription occurred in the S stage of 1-cell zygotes ^14^, and transcription of MuERVL (murine endogenous retrovirus-L) genes was verified via RT-PCR studies^15^. Another study found that transcription was promiscuous across open chromatin regions of the mouse zygote genome, with many transcripts containing intronic and intergenic regions, indicating limited 3’end processing and splicing mechanisms involved, possibly to protect the zygote from precocious gene expression^11^.

Although the understanding of transcriptional events in the zygote is still inconclusive in many species, post-transcriptional regulation of transcripts appears to be ubiquitous. A pivotal aspect of post-transcriptional modification is cleavage and polyadenylation of maternal mRNAs. The choice of PAS in 3’ untranslated regions (3’ UTRs) is thought to influence various UTR binding components that in turn regulate the stability, translation, and degradation of transcripts. Post-transcriptional regulation via alternative poly-A cleavage sites as well as the length of poly-A tail plays vital roles in MZT in many organisms. For example, poly-A tail lengthening promotes global translation in *Drosophila* zygotes ^16^. Removal of de-adenylation components in mouse oocytes leads to developmental arrest in pre-implantation embryos ^17-19^. A recent study has also found extensive remodeling of maternal mRNAs in human embryos via poly-A tail modifications such as changes in poly-A tails and 3’UTR lengths, which are essential for MZT in humans ^20^.

Furthermore, despite the plethora of studies into the embryonic development of mice, little is known about the maternal vs zygote transcriptomic landscape in rats, and how the shared and differing characteristics with mice contribute to early stages of their respective pre-implantation embryos. The rise in application of single-cell transcriptomic studies has opened a front into studying the intricacies of developmental dynamics at the single-cell level. However, many such transcriptomic studies overlook alternative insights that can be made apart from gene expression. In this study, we present orthogonal forms of comparative analysis on the transcriptomic landscape of the early MZT by applying a full-length single-cell RNA sequencing (scRNA-seq) protocol ^21^ to oocytes and zygotes in mice and rats. We identify RNA transcripts that are produced, modified, or degraded in the zygotes. We also reveal mechanisms that are unique and conserved in the two species.

## 2. Materials and Methods

### 2.1 Sample preparation

#### Harvest and preparation of oocytes and zygotes

Mouse and rat oocytes and zygotes were isolated at Horizon Discovery, St. Louis site, operated under approved animal protocols overseen by Horizon Discovery’s Institutional Animal Care and Use Committee (IACUC). C57BL/6N mice and Sprague Dawley rats purchased from Taconic Farms were housed in standard cages and maintained on a 12-h light/dark cycle with *ad libitum* access to food and water. Three- to four-week-old female mice were injected with PMS (5 IU/mouse) 48 hr before hCG (5 IU/mouse) injection, followed by with or without mating to stud males. Fertilized eggs and oocytes were harvested 1 d later, respectively. Four- to five-week-old female rats were injected with 20 units of PMS, which was followed by an injection of 50 units of hCG 48 h later, followed by with or without mating to stud males. Fertilized eggs and oocytes of both species were harvested 1 d later.

#### Construction of scRNA-seq libraries

Full-length double-strand cDNA for each oocyte or zygote was prepared using an sc-RNA-seq libraries SMART-Seq v4 Ultra Low Input RNA Kit (Clontech, Mountain View, CA) following the vendor’s instruction, which was based on the Smartseq2 protocol ^21^. cDNA-seq libraries were constructed using a Nextera® XT DNA Library Preparation Kit (Illumina, Sandiago, CA, Cat. Nos. FC-131-1024, FC-131-1096) following the vendor’s instruction. The libraries were quantified using an Agilent 2100 Bioanalyzer (Agilent Technologies, Santa Clara) and sequenced on an Illumina 2500 machine with 150 bp paired end reads.

### 2.2 Transcriptome mapping and quantification

The mouse genome assembly (GCF_000001635.27) with annotation version GRCm39 and the rat genome assembly (GCF_015227675.2) with annotation version mRatBN7.2 were obtained from NCBI Genbank. Raw sequences were trimmed using Trim_galore ^22^, with parameters length > 35 and q > 10, and subsequently mapped to the respective reference genome using STAR v2.7.10b^23^ with parameters adapted from nf-core/rnaseq workflow settings: **--quantMode TranscriptomeSAM --twopassMode Basic --runRNGseed 0 --outFilterMultimapNmax 20 -- alignSJDBoverhangMin 1 --outSAMattributes NH HI AS NM MD -- quantTranscriptomeBan Singleend --outSAMstrandField intronMotif.** The resulting mappings were quantified using Salmon ^24^ with default settings to obtain estimated expression levels for transcripts in transcript per million (TPM) units. Transcript TPM values were then aggregated via tximport package to obtain estimated gene counts. STAR aligned mappings were also quantified via HTSeq ^25^ to obtain intergenic transcription counts. Quantification of read coverage of nascent transcripts was performed using kallisto^26^/bustools ^27^ through the kb_python package (v0.28.0) with parameters: **–excluded-features pseudogenes** to achieve similar quantification results as that of Salmon quantified expression. More specifically, trimmed reads were aligned to the reference genome with kb_python under the ‘nac’ workflow with default settings and spliced/un-spliced reads were obtained via the kb_python instructions given in ^28^.

### 2.3 Quality control and exploratory analysis

Samples with more than 5% reads mapped to rRNA genes or 25% reads mapped to the mitochondrial genome were removed from further analysis. Furthermore, only genes whose annotated biotypes belong to protein-coding, lncRNA and transcribed pseudogenes were kept for further analysis. Samples were visualized using Uniform Manifold Approximation and Projection (UMAP) based on log transformed DESeq^29^ median normalized counts, outliers were filtered out for further analysis.

### 2.4 Differentially expressed gene (DEG) analysis

Due to the inherent high variability intrinsic to single cell transcriptomic data, to ensure genes had sufficient coverage and replicates for DEG analysis, we only considered genes that were expressed in at least 6 samples. DEG analysis of both mouse and rat samples were carried out with count values from HISAT2^30^/^25^. The raw counts were normalized to Counts per Million (CPM) and log transformed prior to analysis with MAST^31^. To estimate log_2_ fold-change (log_2_FC) values for genes that had no expression in oocytes or zygotes when applying the MAST package, we set parameters: **useContinuousBayes = True.** For each species, genes with an FDR < 0.05 and a foldchange greater than 2 or less than 0.5 were considered differentially expressed.

### 2.5 Estimation of pre-mature mRNA transcripts

Upon estimation of nascent mRNA transcripts via kb_python, the nascent expression matrix and spliced expression matrix was extracted from loom file via scVelo^32^. From henceforth, we shall use the terms un-spliced reads, nascent reads and intronic reads interchangeably to describe the reads that cover intronic portions of genes. To account for bias arising from high technical noise, transcript coverage and putative transcription of unrelated features in the introns of various genes, we adopted a filtering strategy from previous methodologies for RNA ^33,34^, whereby a gene was considered to have pre-mature mRNA expression if it met the following criteria:

- mean read coverage of spliced transcripts across either zygote or oocyte samples > 5.
- spliced transcripts were expressed in 10% of either zygote or oocyte samples.
- mean read coverage of un-spliced transcripts across either zygote or oocyte samples > 0.
- un-spliced transcripts expressed in 10% of either zygote or oocyte samples.
- R^2^ of linear model fit between spliced and un-spliced reads of all samples > 0.1.
- Kendall correlation between spliced and un-spliced reads of all samples > 0.1.
- Ratio of standard deviation of spliced reads over standard deviation of un-spliced reads was between 0.005 and 5.

This filtering criteria was set to ensure that estimation of nascent read coverage truly came from un-spliced intronic portions of transcripts. DEG analysis of un-spliced genes was performed with MAST, with un-spliced reads normalized by sample-wise CPM normalization factors used in the DEG analysis. We define the nascent proportion of a gene as the ratio of the mean spliced read coverage over the mean un-spliced read coverage in a cell. To test differences in nascent proportions (DNP) of a gene between zygotes and oocytes, we applied a Fisher’s exact test.

### 2.6 Differential transcript usage (DTU) analysis

DTU analyses were performed under the guidelines given by ^35^. Specifically, transcript abundances (TPM values) estimated by Salmon^24^ were first filtered using Drimseq^36^ with default parameters. The filtered data was then analyzed using DEXseq^37^ with default settings. Instead of exons in the original applications ^37^, each transcript of a gene was regarded as an exon in our case. The DEXseq results for each gene were aggregated and further analyzed using stageR^38^ to determine if it exhibited DTU while overall false discovery rate (FDR) was controlled on the gene level.

### 2.7 Alternative polyadenylation analysis

STAR aligned BAM files were first filtered using Samtools v1.19^39^ to only include concordantly and uniquely mapped reads. The resulting filtered bam files were then transformed into BEDGRAPH format using Samtools depth tool along with custom scripts^40^. Primary lists of mouse and rat 3’UTRs were obtained by extracting 3’UTRs from the NCBI annotation files via custom scripts and only unique 3’UTRs were kept. Though most UTRs were composed of a single continuous segment, there was still a portion of UTRs that contained introns. In these cases, we treated split the intron containing UTRs and treated each segment as individual 3’UTRs. We identified genes with differential alternative polyadenylation (DAP) in each species using DaPars2 ^41,42^, which computed a Percentage of Distal poly-A site Usage Index (PDUI) by de novo estimating putative polyadenylation sites (PAS) via read coverage change on the lists of 3’UTRs. To ensure a more accurate DAP analysis, we modified DaPars2 as follows:

1. Begin search for putative PAS 25 bp downstream instead of the default 150 bp downstream of 3’UTR start.
2. Furthermore, instead of using read depth of each sample for normalization of each sample’s read depth, we used DESeq2-estimated size factor of each sample’s read counts as normalization factor.
3. Modified DaPars2 to allow output of distal and proximal PAS coverages in addition to PDUI units for each UTR.

Moreover, the threshold for coverage was set to 0, thus, any read coverage was counted in the estimation of alternative polyadenylation. To ensure robust estimated 3’UTR coverage prior to testing, we filtered out genes that satisfied the following criteria:

1. Average estimated reads less than 10 reads (STAR-Salmon estimated counts) detected in across all samples,
2. Genes with non-NA 3’UTRs coverage in < 3 samples in both conditions.
3. Average 3’UTR coverage depth > 2 for either long or short 3’UTR for both conditions.

UTR regions were further filtered to only retain regions with existence of common PAS signals in nearby regions of the estimated PAS (a PAS motif must be found within 80bp upstream and 120bp downstream of the estimated PAS to be considered valid for testing). We then performed Fisher’s exact test to evaluate the differences in distal vs proximal PAS preference between oocytes and zygotes using mean distal/proximal coverage in oocyte and zygote samples. Differences in mean PDUI between oocytes and zygotes were also calculated. To aggregate the results to gene level (many genes have multiple 3’UTRs), for each gene, we combined p-values of all its viable UTRs using the ‘**simes**’ method implemented by the **metapod** package^43^. The PDUI value of a gene with multiple significant DAP 3’UTRs was defined as the PDUI of the 3’UTR with the largest absolute PDUI value. The resulting p-values were then controlled for FDR with the qvalue package in R. Finally, genes with UTRs that had an absolute mean expression difference > 0.2 and a FDR < 0.05 were considered to have significant differential alternative polyadenylation. Gene-body coverage in each sample was normalized by the BEDGRAPH coverage as aforementioned, and the average was taken over the oocyte or zygote samples. The normalized mean 3’UTR coverage plots for a gene was obtained via averaging coverage depth across all samples of each condition via custom scripts and visualized using the trackViewer R package^44^.

### 2.8 Pathway and GO term enrichment analysis

The ClusterProfiler^45^ R package was used to perform gene set enrichment analysis (GSEA) for results of differential expression and over-representation analysis (ORA) for results of DNP, DTU, DAP and orthology analysis. Gene sets/pathways combined from KEGG ^46,47^, Gene Ontology Biological Processes (GO BP)^48^, Reactome ^49^ and Wikipathways^50^ for were used for analysis.

### 2.9 Analysis of orthologs between mouse and rat

Orthologs between mice and rats were obtained from the MGI database^51^. Specifically, rat and mouse genes belonging to the same homology class were considered as orthologs. ORA for mouse-based gene sets and pathways was performed on orthologous genes that had an absolute foldchange > 1.25 and FDR < 0.05 in both species, with all orthologous genes that were tested for differential expression in both species as the gene background. Visualizations of gene networks were generated using clusterProfiler^45^. All intersections of genes with DNP, DTU and DAP genes were performed with the respective results obtained in the prior analysis. All gene coverage plots were generated via the trackViewer^44^ package.

## 3. Results

### 3.1 Oocytes and zygotes of both mice and rats display distinct transcriptomic patterns

We sequenced the transcriptomes of a total of 17 mouse oocytes, 17 mouse zygotes, 15 rat oocytes, and 16 rat zygotes, with an average of around 10 million paired-reads/cell (Supplementary Table S1). The vast majority (>90%) of the reads could be mapped to the respective genomes for most (53% to 81%) samples (Supplementary Table S1). However, 3 mouse oocytes, 1 rat oocyte and 5 rat zygotes exhibited high levels of reads mapping to mitochondrial or ribosomal genes, we thus excluded them from further analyses (Supplementary Table S1) (Materials and Methods). This left us with 31 mouse samples (14 oocytes / 17 zygotes) and 25 rat samples (14 oocytes / 11 zygotes). Interestingly, the reads from mouse samples, particularly, from mouse oocytes, were strongly biased to the 3’ ends of CDSs of genes, while the bias was not seen in rat oocytes and zygotes (Supplementary Figure S1A). We detected transcripts for an average of 14978 and 15604 annotated genes in mouse oocytes and zygotes, respectively, and for an average of 12,829 and 13,468 annotated genes in rat oocytes and zygotes, respectively (Supplementary Figure S1B). Though not significant, there appear to be more genes expressed in zygotes than in oocytes of both species (Supplementary Figure S1B, Wilcoxon rank-sum test, p-value < 0.1). Furthermore, oocytes and zygotes of both species are clearly separated by estimated gene expression levels in UMAP embeddings (Figures 1A, 1B), indicating that oocytes and zygotes of both mice and rats contain distinct profiles of gene expression.

**Figure 1.**
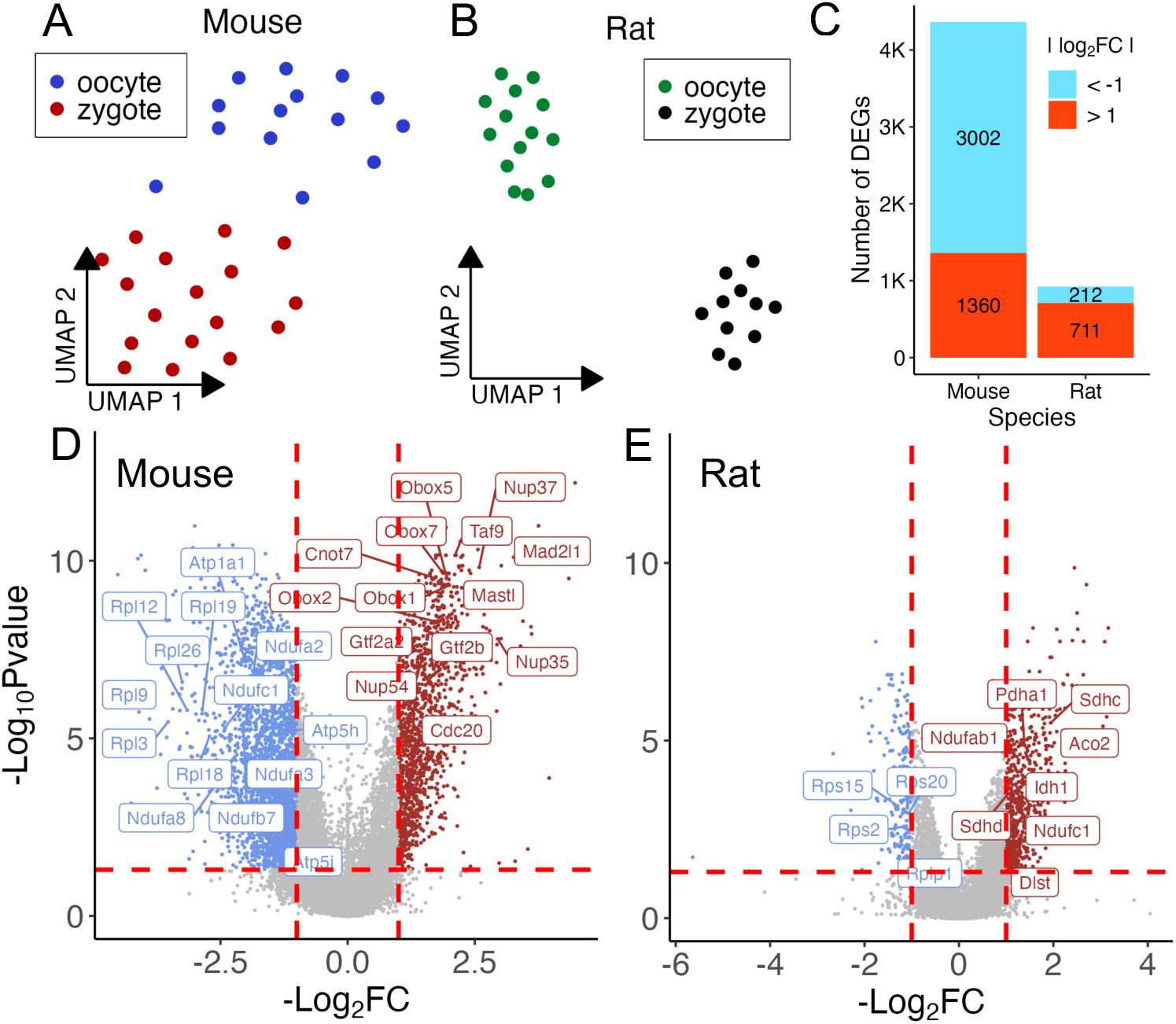
Oocytes and zygotes of both species contains distinct sets of transcripts. **A.** UMAP visualization of single mouse oocytes and zygotes based on expression levels of genes. **B.** UMAP visualization of single rat oocytes and zygotes. **C.** Number of up- or down-regulated DEGs in mouse and rat zygotes relative to respective oocytes. **D, E.** Volcano plots of –log_10_Pvalues vs log_2_FC of mouse (D) and rat (E) genes. Genes marked red were significantly upregulated (log_2_FC > 1 and FDR < 0.05), while those marked blue were significantly downregulated (log_2_FC < -1 and FDR < 0.05) in zygotes relative to oocytes.

### 3.2 ZGA in rats may begin earlier than previously believed

We first compared expression levels of genes in zygotes relative to those in oocytes of each species. We found 4,362 DEGs in mice, of which 1,360 or 29.7% were upregulated while a larger portion, 3,002 or 70.3% of DEGs, were downregulated (Figure 1C, Supplementary Table S2). In contrast, we found only 923 DEGs in rat, of which a majority, 711 or 74.5% of DEGs, were upregulated while the remaining, 212 or 25.5%, were downregulated (Figure 1C, Supplementary Table S3). Since zygotes of both species were analyzed prior to the first cell division, the elevated transcriptional activities of the 1,360 upregulated DEGs in mice may reflect initial transcriptional events in mid-1cell stage zygotes reported in early studies^11,52^. Despite rat zygotes exhibiting a relatively quiescent differential expression landscape compared to mice, the large proportion of upregulated DEGs in rats suggests an early wave of transcriptional activation also occur in rat zygotes, which was previously believed to happen in later stage of embryogenesis ^12^.

### 3.3 Distinct pathways are up- or down-regulated in both mouse and rat zygotes

GSEA analysis found downregulation of gene sets/pathways including cytoplasmic translation, rRNA processing, nonsense-mediated decay, TCA cycle and chromatin modifying enzymes, etc. (Figures 1D, 2A, Supplementary Table S4). For example, ribosomal protein subunits encoding genes such as *Rpl*3/9/12/18/19/26 ^53^ were downregulated in mouse zygotes (Figure 1D), a result that is in excellent agreement with the earlier findings ^54^. Upregulated gene sets/pathways included mRNA transport, cell cycle, transcription, DNA replication, etc. (Figures 1D, 2A, Supplementary Table S4). Surprisingly, genes involved in energy production exhibit both upregulation and downregulation. For instance, genes that function in the TCA cycle such as *Ndufa2/Nudfb7/Ndufa8/Ndufc1* ^55^ and genes involved in energy production such as *Atpa1a/Atp5h* ^56^ showed decreased transcript levels (Figure 1D). On the other hand, we found upregulation of genes *Pdhb, Pdp1, Pdhx* and *Dlat*, which encode components of the pyruvate dehydrogenase complex responsible for acetyl-CoA biosynthesis from pyruvate ^57^(Figures 1D). Interestingly, previous early mouse embryos studies on the role of the TCA cycle in ZGA found that despite low metabolic requirements in early embryos, pyruvate was responsible for the nuclear localization of several TCA enzymes that might contribute to epigenetic regulations that were essential for major ZGA activation in the 2-cell stage^58^, providing a possible explanation for the dichotomy seen here. RNA polymerase II subunit genes *Taf*6/9 ^59,60^, cell cycle genes *Cdc20, Mastl,* and *Mad2l1* ^61-63^, and general transcription factor II genes *Gtf2a2/2b*^64^ also exhibited upregulation in mouse zygotes (Figure 1D). We also observe nucleoporin (Nup) encoding genes *Nup35/37/54*, and the nuclear transport factor 2-like export factor ^65^ encoding gene *Nxt1* were upregulated, suggesting increased transport of mRNA and proteins between the nucleus and cytoplasm in mouse zygotes, in line with recent findings that knockdown of the nucleoporin gene *Nup37* led to reduced blastocyte formation rates ^66^ and that massive remodeling of the nuclear envelope occurred during minor zygotic activation in bovine pre-implantation embryos^67^. Moreover, we found genes encoding essential ZGA transcription factors *Obox1/2/5/7* ^68^ were all upregulated in mouse zygotes (Figure 1D), suggesting that the elevation of transcription levels of these genes during minor ZGA may also be at least partially transcriptional and not solely of maternal origin as reported by a recent study ^68^.

**Figure 2.**
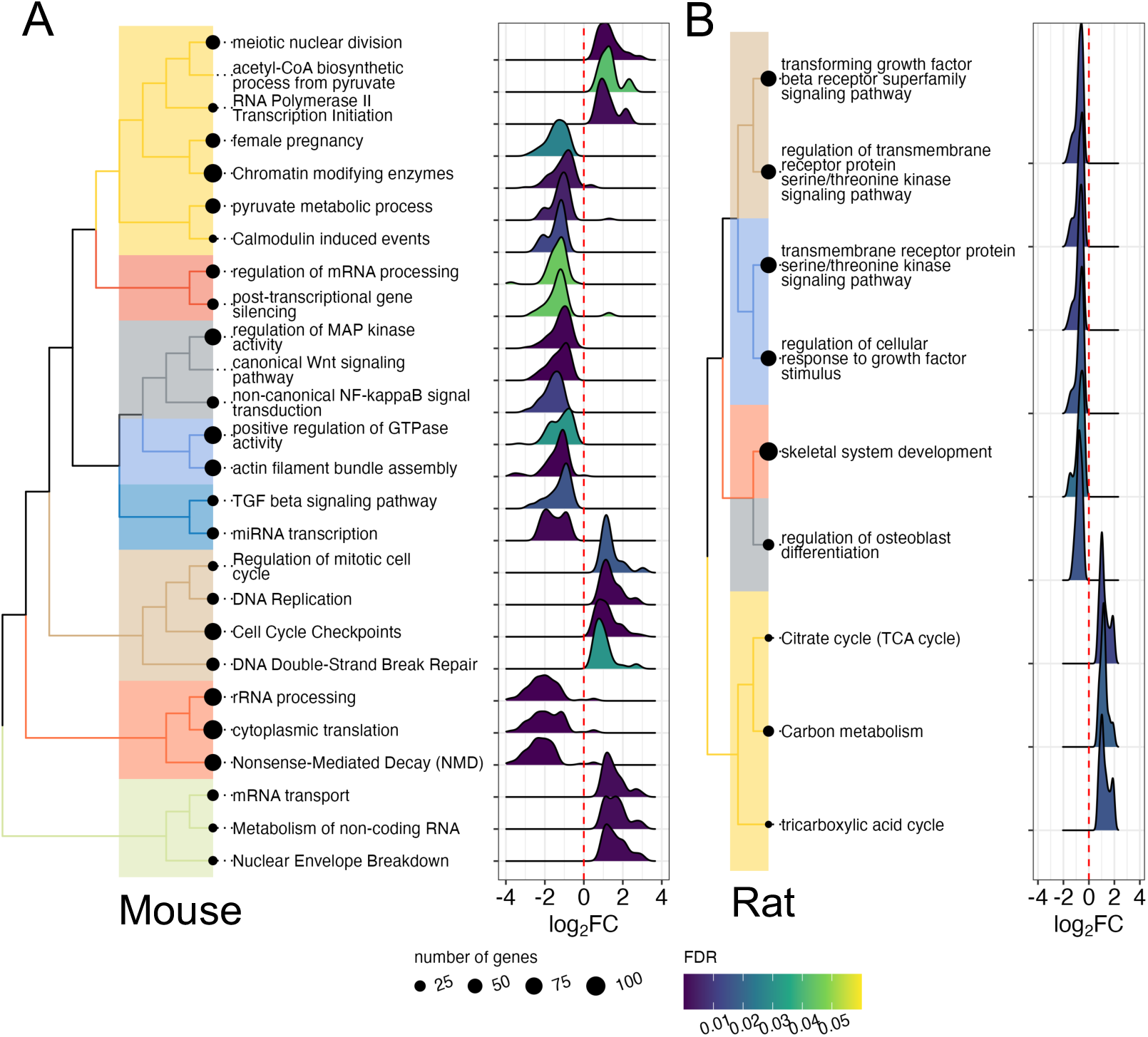
**A,B.** Enriched GO BP terms, Wikipathways, KEGG Pathways and Reactome at FDR < 0.05 in mouse (A) and rat (B). Significant pathways and gene sets are clustered into similar groups based on Jaccard’s similarity for ease of viewing, density plots for each gene set/pathway show distribution of the log_2_FC of the genes in the indicated GO BP or pathways.

Downregulated gene sets/pathways in rat zygotes involve growth factor stimulation such as the transforming growth factor beta receptor superfamily signaling pathway (Figure 1E, 2B, Supplementary Table S5), which has been known to regulate cell growth, differentiation, and other cellular functions^69^. Upregulated genes in rat zygotes significantly enriched for carbon metabolism pathways and the TCA cycle, etc (Figures 1E, 2B, Supplementary Table S5). For instance, transcription of the following genes involved in the TCA cycle were all upregulated in rat zygotes: *Aco2* (aconitase 2) ^70^, *Pdha1*(pyruvate dehydrogenase component) ^57^, *Idh1* (pyruvate dehydrogenase) ^71^, *Sdhc/d* (succinate dehydrogenase subunits) ^72^, and *Dlst* (dihydrolipoamide S-succinyltransferase) ^73^(Figure 1E). This result contrasts with that seen in mouse zygotes where some energy production genes of the TCA cycle were downregulated while others were upregulated (Figures 1D, 2A). Coupled with the aforementioned studies showing that several TCA cycle enzymes were transiently localized in the nucleus of mammalian embryos and essential for major ZGA ^58,74^, the shared increased expression of the said enzymes in both species suggests that these enzymes may be regulated at the transcriptional level during minor ZGA. Taken together, these results once more suggest that ZGA in the rat might begin earlier than previously believed.

### 3.4 Pre-mature mRNA transcription is elevated in zygotes of both species

It has been reported that early transcriptional events in the zygote produced pre-mature (nascent) RNAs that originated from intronic as well as intergenic regions^75^. We thus estimated nascent (un-spliced) and mature (spliced) transcripts in the samples of both species (Materials and Methods). The estimated spliced transcript read counts for each sample strongly correlated with our estimated read counts used for differential gene expression (Supplementary Figure S2A). We detected a total of 26,327 and 20,745 genes with read coverage in the mouse and rat oocytes and zygotes, of which 2989 (11.4%) and 2892 (13.9%) had nascent reads respectively after filtering (Materials and Methods, Supplementary Table S6, S7, Supplementary Figure S2B). In both species, zygotes contained higher proportions of nascent reads than oocytes (Figure 3B). Previous studies also reported a higher percentage of intergenic read coverage in zygotes ^75^, thus we also quantified reads mapped to 1 kb binned intergenic regions. Zygotes in both species had significantly greater number of intergenic bins with at least 2 read coverage than oocytes, either across all intergenic bins or those that are at least 10kb from any gene (Figure 3C). These results corroborate previous findings of increased nascent and intergenic transcription in mouse zygotes, while also suggesting a similar occurrence in rat zygotes. We then performed a differential expression analysis of genes with nascent transcripts in oocytes and zygotes (Materials and Methods) to obtain fold changes of nascent transcripts. We found that log_2_FC values of nascent transcripts and spliced transcripts were strongly correlated (Figure 3C), indicating that nascent transcripts mostly changed in the same ways as their spliced counterparts.

**Figure 3.**
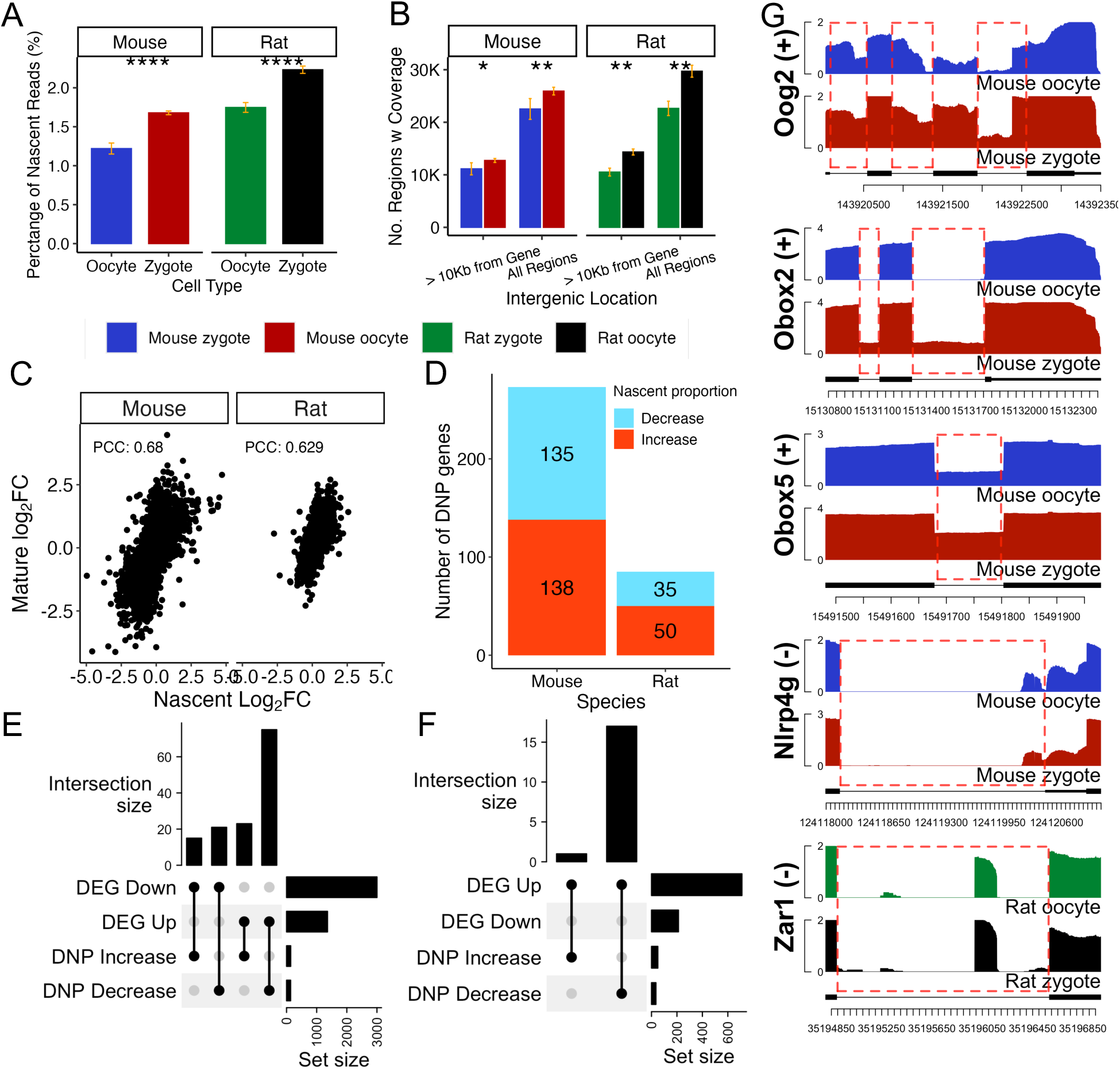
Expression of pre-mature mRNA transcripts in oocytes and zygotes of both species. **A.** Average percentage of total un-spliced reads out of all reads (spliced and un-spliced) in oocytes and zygotes of mice and rats. **B.** Number of all and far 1kb intergenic regions with reads in oocytes and zygotes of the two species. Far regions are >10kb downstream or upstream of any gene. **C.** Scatter plots of MAST estimated log_2_FC values of un-spliced reads and log_2_FC of spliced reads for all genes that have un-spliced reads in each species. Pearson’s correlation coefficient (PPC) between the values is shown for each species. **D.** Number of genes with differential nascent proportion (DNP) for each species, which are up- or down-regulated in zygotes. **E, F.** Upset plots of increased/decreased DNP genes and upregulated/downregulated DEGs in mice (E) and rats (F). **G.** Average intronic coverage for *Oog2, Obox2, Obox5, Nlrp4g* and *Zar1* that were called DNP, in oocytes and zygotes, y-axis is log_10_ transformed read coverage, red boxes highlight regions with intronic coverage. * p < 0.05, ** p<0.01 and *** p<0.005, Wilcoxon rank-sum test.

To identify genes that exhibited significant changes in the proportion of nascent transcripts, we performed a DNP analysis between oocytes and zygotes in both species (Materials and Methods). We found that mouse had similar number of genes with decreased nascent proportions than with increased nascent proportions (135 vs 138) (Supplementary Table S6), while rat had more genes with increased nascent proportions than with decreased nascent proportions (50 vs 35) (Figure 3D) (Supplementary Table S7). Comparisons of DEGs and DNP genes in both species found only small overlaps between DEGs and DNPs (Figure 3E, 3F). However, in both species, the largest overlap was between upregulated DEGs and genes with decreased nascent proportions (Figure 3E, 3F). Although this was expected in rats due to a greater number of upregulated DEGs than downregulated DEGs (Figure 2C), it was surprising in mice, since mouse had many more down-regulated DEGs than upregulated DEGs (Figure 3E). This result suggests that some of upregulated DEGs in mice could be due in part to more mature transcripts in the zygote cytoplasm that may have been stored as unprocessed transcripts in the oocyte nucleus. ORA of these DNP genes in both species yielded no significant gene sets/pathways when corrected for FDR suggesting that most of the nascent transcription may indeed be pervasive and in-discriminant ^75^. However, we did find changes in genes that played important roles in either oocytes or zygotes. For instance, the nascent proportion of the *Oog1/2* genes increased in mouse zygotes (Figure 3G), which is known to be required for oocyte development ^76,77^. The nascent proportion of the *Obox* genes increased in mouse zygotes (Figure 3G), consistent with our earlier observation that the *Obox* genes were upregulated in the zygotes as a part of the minor ZGA process, albeit in unprocessed transcript form. The *Nlrp4b/g* genes exhibited decreased nascent proportions in mouse zygotes (Figure 3G), in agreement with the previous reports that the *Nlrp* transcripts were maternally derived and played essential roles in the zygotic genome ^78,79^. Although DNP genes in rats were less interesting, we found the *Zar1* gene to exhibit decreased nascent transcript proportion (Figure 3G), consistent with the previous report that Zar1 played essential roles in zygotic genome activation by regulating maternal derived RNAs ^80^.

### 3.5 Differential splicing might play a role in MZT in both mice and rats

To uncover genes with splicing changes and isoform switches during MZT, we performed DTU analysis (Materials and Methods) on genes between oocytes and zygotes of both species. We found that 2,869 genes exhibited significant DTU between mouse oocytes and zygotes, while only 601 genes showed significant DTU between rat oocytes and zygotes (Figure 4A, Supplementary Table S8, S9), once more suggesting a much more dynamic transcriptional landscape in the mouse zygotes than in rat zygotes. In both species, most significant differential transcript usage of a gene can be attributed to the significant change of no more than two isoforms of that gene. (Supplementary Figure S3A). Only small portions of significant DEGs overlapped genes with significant DTU genes in both species (Figure 4B), suggesting largely decoupled sets of transcriptionally regulated and post-transcriptional modified targets in the zygote. Interestingly, the number (388) of DTU genes in mice that also are upregulated DEGs is close to the number (415) of DTU genes that also are downregulated DEGs, despite many more downregulated DEGs in mice (Figure 4B). Genes called for DTU in mice significantly enriched for processes such as cell cycle, epigenetic regulation of gene expression and chromatin organization/remodeling (Figure 4C, Supplementary Table S10). Though DTU genes in rats did not significantly enrich for any gene sets or pathways (FDR < 0.05) (Figure 4D, Supplementary Table S11), top ranked gene sets (by p-values, without multiple hypotheses FDR correction) for rats also include epigenetic regulation of gene expression and chromatin organization/remodeling (Figures 4D, Supplementary Table S11). It has been shown that the chromatin undergoes drastic changes during MZT including redistribution of histone H2A among other epigenetic makers, and thus, the control of H2A is vital to ensure proper embryonic development ^81^. Previous studies also suggest that both accumulation and absence of H2A may result in embryonic arrest^82,83^. Moreover, we found *Bap1* and *Usp3* exhibiting DTU in mice (Figures 5A, 5B), and *Usp16* and *Rnf2* showing DTU in rats (Figures 5C, 5D). These results are consistent with earlier findings that *Usp3* and *Bap1* deubiquitinate H2AK119ub1 ^84,85^, *Usp16* is the dominant deubiquitinase of H2AK119ub1 in oocytes^83^, and Rnf2 monoubiquitinates H2A histone ^86^. Taken together, our results suggest that alternative splicing might serves role in regulating genes involved in epigenetic regulation in the ZGA of both species.

**Figure 4.**
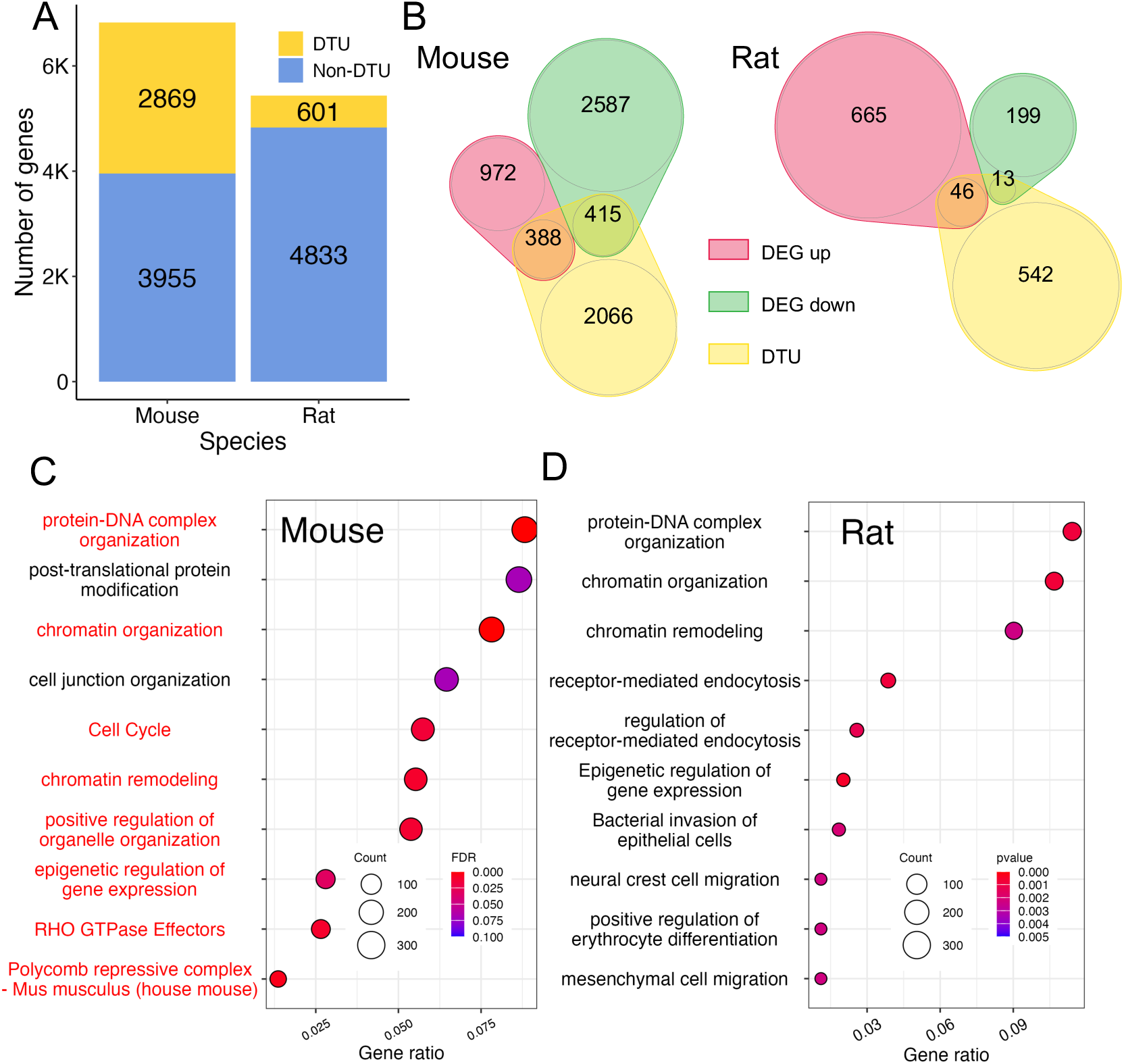
Differential splicing of transcripts between oocytes and zygotes of mice and rats. **A.** Number of transcribed genes with and without DTU (FDR < 0.05) in each species. **C.** Venn diagram of DTU genes and upregulated or downregulated DEGs in each species. **D.** GO BP enrichment for mouse genes with DTU (top 10 terms are shown, names marked in red are significant, FDR < 0.05). **E.** Go BP enrichment for rat genes with DTU (top 10 terms are shown). Gene ratio is the ratio of genes in a GO BP term that were called DTU over all genes in that term.

**Figure 5.**
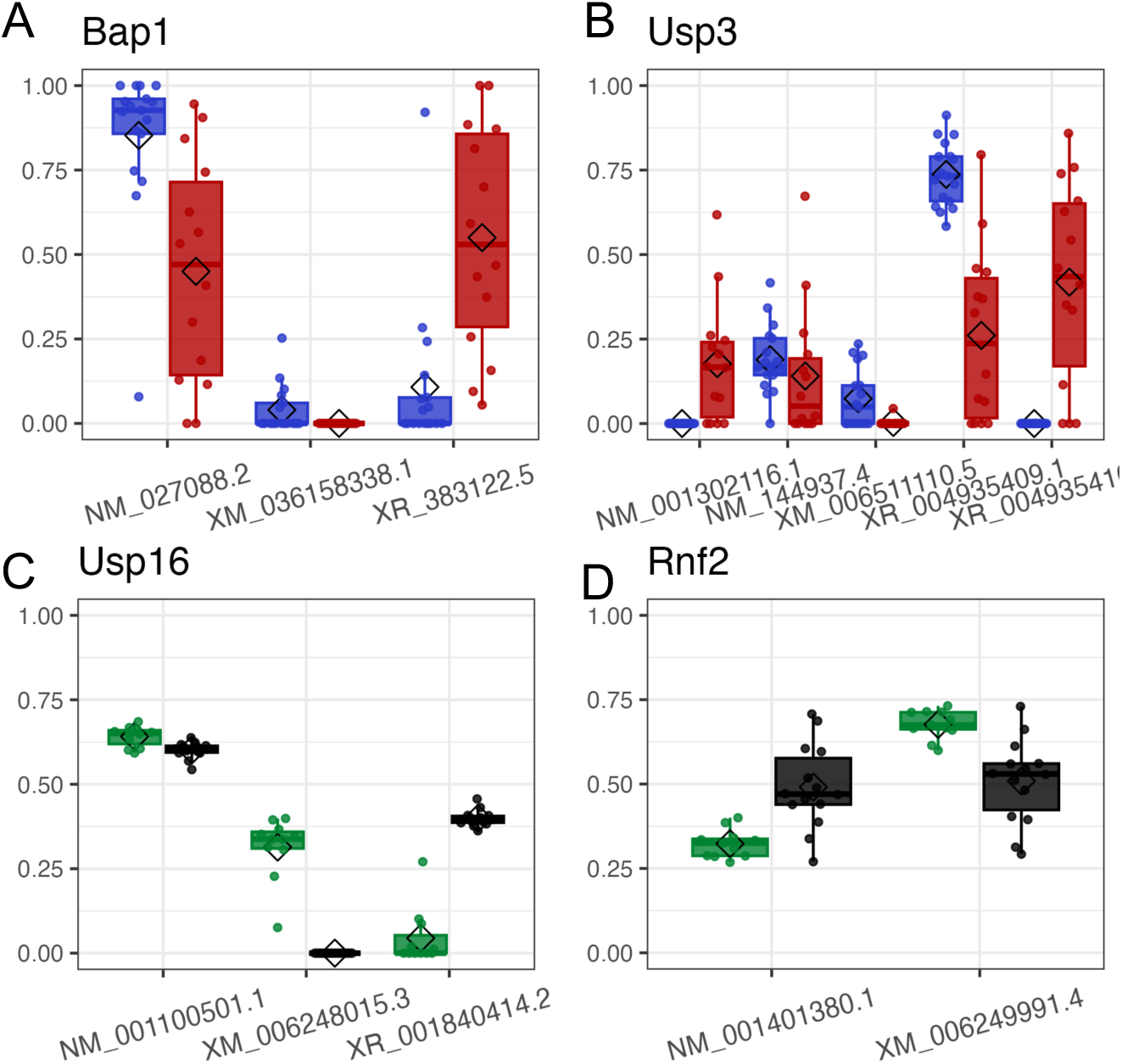
**A-D Comparison of** proportions of isoforms of *Bap1*(**A***), Usp3* (**B**), *Usp16* (**C**) and *Rnf2* (**D**) between oocytes and zygotes of mice and rats.

### 3.6 Gene transcripts in oocytes and zygotes undergo distinct 3’UTR polyadenylations in both mice and rats

Post-transcriptional modifications to mRNA molecules such as alternative use of PAS in 3’UTRs, play a critical role in the regulation of mRNA stability and translation ^87,88^. Thus, we analyzed differences in 3’UTR usage of transcribed genes between oocytes and zygotes in both species by calculating a PDUI (Percentage of Distal poly-A site Usage Index) value ^41,89^ for each expressed gene (Materials and Methods). Oocytes and zygotes in each species could be clearly differentiated based on their PDUI values in UMAP embeddings (Figures 6A, 6B), indicating that genes expressed in oocytes and zygotes underwent distinct 3’UTR polyadenylations. We then sought to uncover significant differences in 3’UTR usage between oocytes and zygotes (Materials and Methods). In mice, we found that 580 genes exhibited significant DAP between oocytes and zygotes; of which, 420 (72.4%) showed increases in proximal PAS usage (preference for shorter 3’UTRs), while the remaining 160 (27.6%) showed preference for distal PAS (preference for longer 3’UTRs) (Figure 6C, Supplementary Table S12). In rats, we found that 121 genes showed significant DAP between oocytes and zygotes; of which, 66 (54.5%) had increased preference for distal PAS, while the remaining 55 (45.5%) showed increased preference for proximal PAS (Figure 6C, Supplementary Table S13). DAP genes with shortened 3’UTRs in mice are enriched for mitotic cell cycle transition as well as meiotic division (Figure 6D, Supplementary Table S14), while no significant gene sets/pathways were enriched for genes with lengthened 3’UTRs (Supplementary Table S15). In rats, only actin filament organization was significantly enriched for the DAP genes with lengthened 3’UTRs after FDR correction (Supplementary table S16), though top ranked (by p-value) gene sets/pathways enriched by significant DAP genes in rats include positive regulation of G2/M transition of mitotic cell for both lengthened and shortened 3’UTR genes (Supplementary table S16, S17).

**Figure 6.**
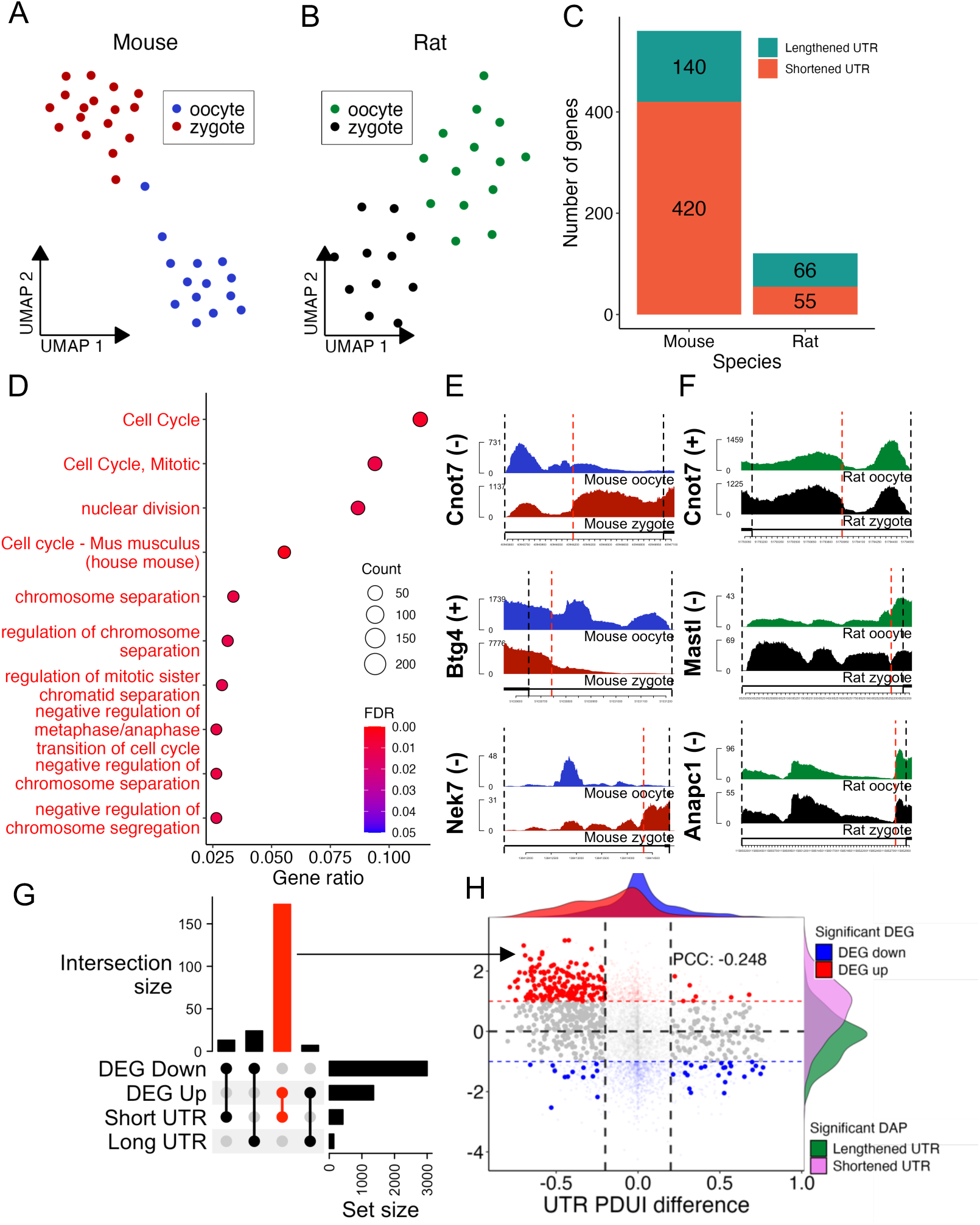
Differential alternative polyadenylation of transcripts between oocytes and zygotes of mice and rats. **A, B** UMAP display of mouse (A) and rat (B) oocytes and zygotes based on PDUI values of genes. **C**. Number of genes with lengthened and shortened UTRs in zygotes relative to in oocytes of mice and rats. **D.** Significantly enriched GO BP terms for DAP genes in mice (FDR < 0.05). **E.** Reads coverage on the 3’UTRs of mouse genes *Cnot7, Btg4* and *Nek7*. **F.** Read coverage on the 3’UTRs of rat genes *Cnot6l, Mastl* and *Anapc1.* In both H) and I), red lines mark the predicted proximal PAS. **G.** Upset plot of DAP genes and DEGs in mice. **H.** Log_2_FC expression vs PDUI difference for all genes that were analyzed for DAP; low transparency points are genes not significant for DAP, red/blue genes are significantly up/down regulated respectively; pink and green density plots show log_2_FC distribution for significant DAP genes with lengthened and shortened UTRs; red and blue density plots show PDUI changes for significantly up/down regulated DEGs.

Upon closer inspection, many of the DAP genes in mice or rats are known to play various roles in the MZT. We describe some examples of these genes below. The first set are involved in deadenylation and readenylation of mRNAs. We found that *Btg4* exhibited DAP with shortened 3’UTRs in mice (Figure 6E) while *Cnot7* exhibited shortened 3’UTRs in both mice and rats. It has been shown that CNOT7 along with CNOT6/6l/8 form the CCR4-NOT complex that controls deadenylation of mRNAs ^19,90-92^. It has also been recently reported that BTG4 functions as a mediator of deadenylation, playing a critical role in the production of substrates for maternal RNA re-adenylation during MZT in humans^7,17,18^. Our results suggest a putative mechanism by which these deadenylation related genes switch to shorter 3’UTRs in zygotes to increase translational efficiency, which in turn drives further de-adenylation (and subsequent readenylation) and shortening of 3’UTRs of other genes, thus conferring higher translational efficiency of genes crucial for zygote development.

Many genes are implicated in the regulation of mitotic progression and cell cycle, particularly in mice. For instance, we found that *Nek7* had DAP with shortened 3’UTR in mouse zygotes (Figure 6E). An early study has shown that reduction in *Nek7* activity can cause cells to arrest in metaphase of mitosis ^93^. It is likely that *Nek7* serves a similar function in meiosis through an inhibitory mechanism due to a longer 3’UTR in MII oocytes. Interestingly, unlike the case in mice, several mitosis related DAP genes in rats exhibited lengthened 3’UTR usage. The *Mastl* gene, which encodes an essential regulator of cell cycle control ^61^, exhibited increased usage of longer 3’UTR transcripts in rat zygotes (Figure 6F). Similarly, *Anapc1* exhibited increased usage of longer 3’UTRs (Figure 6F). *Anapc1* encodes a subunit of the Anaphase Promoting Complex/Cyclosome (APC/C), a key ubiquitin-ligase in mitosis and it has been shown that delay in the APC/C activation extends mitosis in mouse embryos ^94^. Though not significant in mice, a similar change in 3’UTR usage can be seen for mice *Anapc1* (Supplementary Figure S3B). Taken together, our results suggest a putative regulatory mechanism of *Anapc1* via 3’UTR extension, which may in turn limit translational efficiency and cause delays in subsequent mitosis ^95^. These results suggest changes in 3’UTR usage may serve as a post-transcriptional regulatory mechanism affecting mRNA stability and protein translation in rat and mouse zygotes during the first mitotic progression.

We then investigated the relationships between DEGs and DAP genes. In mice, a considerable number of DAP genes were also DEGs. Specifically, of the mouse DAP genes with shortened 3’UTRs, 173 were upregulated DEGs, but only 13 were downregulated DEGs, despite fewer upregulated DEGs than downregulated DEGs (Figure 1C). For DAP genes with lengthened 3’UTRs in mice, 7 were upregulated DEGs, and 24 was downregulated DEGs (Figure 6D). These results suggest that DAP genes in mice with shortened 3’UTRs tend to be upregulated DEGs (Figure 6G). In addition, we analyzed the log_2_FC changes of all genes analyzed for DAP (Figure 6H) and found that though genes with shortened 3’UTRs tended to be upregulated (skewed towards log_2_FC > 0), though there appeared to be no correlation (PCC: -0.248) between the magnitude of expression change and changes of 3’UTR usage (PDUI difference). We also found that significant DAP genes with lengthened 3’UTRs showed no apparent association with upregulation/downregulation (Figure 6H). In rats, only a few DAP genes were also DEGs, with no correlation between the numbers of DEGs and DAPs (Supplementary Figure S3C), thus no confident conclusion could be drawn.

### 3.7 Most orthologous genes in mice and rats have similar while a small portion have opposite transcriptional patterns

We sought to investigate to what extent DEG, DNP, DTU and DAP genes in mice and rats are conserved based on their orthologous relations. Surprisingly, even though the two species share many orthologous genes (14,248 genes with expression in both species), only about 7.7% (1091) were found to be significantly differentially expressed (FDR < 0.05, |log_2_FC| > log_2_1.25) in zygotes relative to in oocytes in both species. However, most (753, or 69%) of the orthologous DEGs showed similar tendency of change (341 upregulated and 412 downregulated) (Figure 7A), while the rest small portion (338 or 31%) exhibited opposite changes, i.e., 101 upregulated DEGs in mice were downregulated DEGs in rats, and 237 downregulated DEGs in mice were upregulated DEGs in rats (Figure 7A). These results suggest that most of the orthologous genes that exhibit changes in both species behave similarly in terms of transcriptional change. ORA of these orthologous DEGs reveal enrichment for multiple gene sets/pathways that may be crucial to MZT in both species (Figures 7B, C). Orthologous DEGs that were mostly upregulated in both species were enriched for transport of transcripts and cell cycle, while DEGs that were downregulated in both species were enriched for chromatin remodeling and Nonsense-mediated decay. Orthologous DEGs that were upregulated in rats but downregulated in mice were enriched for aerobic respiration, RNA splicing, translational initiation and rRNA processing (Figures 7B, C).

**Figure 7.**
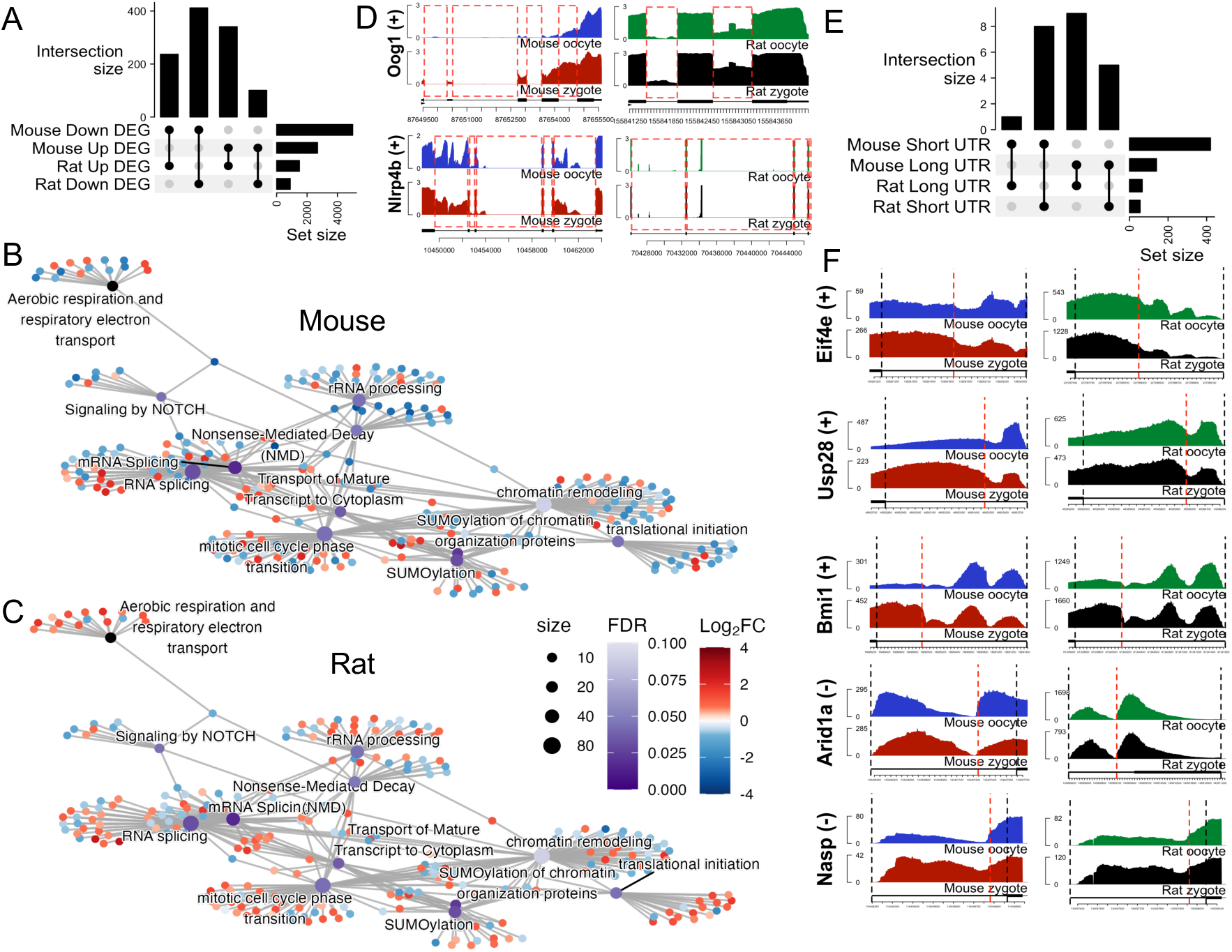
Orthologous genes in mice and rats may have similar or opposite transcriptional patterns. **A.** Upset plot of upregulated and downregulated orthologous DEGs in zygotes relative to oocytes of mice and rats. **B, C.** Enriched GO BP terms for orthologous DEGs in mice (B) and rats (C). **D.** Read coverages of orthologous DNP genes *Oog1* and *Nlrp4b* in oocytes and zygotes of mice and rats. **E.** Upset plot of orthologous DAP genes in mice and rats. **F. 3’**UTR coverage of orthologous DAP genes *Eif4e, Usp28, Bmi1, Arid1a,* and *Nasp* in mice and rats.

Only 6 orthologous genes with DNPs in both mice and rats exhibited similar changes in nascent transcript proportions (Supplementary Figure S4A). Nonetheless, some may play essential roles in MZT. For instance, the *Oog1* gene encoding oogenesin exhibited increased nascent proportions in both mouse and rat zygotes (Figure 7D). It has been shown that oogenesin plays critical roles in zygotic transcription^76^. Moreover, *Nlrp4b* showed decreased nascent proportion in both mouse and rat zygotes (Figure 7D). Studies have shown that *Nlrp4a-g* may play important roles in early pre-implantation embryos, with high expression of NLRP4G and NLRP4E in both oocytes and zygotes ^78,79^.

There were 252 orthologous genes with DTU in both mice and rats (Supplementary Figure S4B), though no gene sets/pathways were significantly enriched. Finally, a total of 23 orthologous genes had DAP in both species, with either lengthened or shortened 3’UTR in both species mostly in accordance (Figure 7E). Close inspection found many interesting orthologous genes exhibiting similar significant changes in 3’UTR usage. For instance, *Eif4e* (Figure 7F), which encodes a rate-limiting regulator of translation of ^96,97^, exhibited preference for shortened 3’UTR in both rat and mouse zygotes (Figure 7F), implying higher translational efficiency of the transcripts to prepare the zygotes for higher levels of protein production in the later stages. It has also been shown that Eif4e interacts with CNOT7 via BTG4, thereby expediting maternal mRNA degradation during MZT ^17,18^. Moreover, both *Usp28* and *Bmi1* exhibited shortened 3’UTRs in both mouse and rat zygotes (Figure 7F). Though the products of both genes participate in the regulation of histone H2A, it has been shown that Usp28 is a deubiquitinase of H2A^98^, while Bmi1 forms the core protein of the Polycomb Repressive Complex 1 (PRC1), which mono-ubiquitinates H2A histones^99^. This result, along with the DTU changes mentioned prior, suggests post-transcriptional splicing and polyadenylation both participate in regulating chromatin states for ZGA in mice and rats. In addition, both *Arid1a* and *Nasp* exhibited lengthened 3’UTRs in both mouse and rat zygotes (Figure 7F). *Arid1a* encodes a subunit of chromatin remodeling complexes, and its expression in mouse embryos has been found to accumulate during G0 phase and vanish by mitosis ^100^. The *Nasp* gene encodes nuclear autoantigenic sperm protein (NASP), which has been found to be present in the oocyte and zygotes of mice^101^, is regulated during cell cycle progression^102^ and blocks G_1_/S progression when overexpressed^103^. The lengthened 3’UTR of *Arid1a* and *Nasp* transcripts in both mouse and rat zygotes may represent a snapshot in the first cell cycle of the embryo, implying a post-transcriptional regulating mechanism through extension of 3’UTR tails^100^.

## 4. Discussion

Our analysis of DEGs between oocytes and zygotes of mice and rats revealed new different facets of the underlying workings of their early MZT. Most prominently, we found decreases in ribosomal protein and nonsense-mediated mRNA decay related transcripts in zygotes of both species, though to a lesser degree in rats. We also found significant increases in transcripts of genes related to the cell cycle, mRNA transport and transcription initiation in mouse zygotes. Although most transcripts in mouse zygotes undergo degradation upon fertilization, there are clearly newly produced transcripts. Moreover, the increased transcripts of nucleoporin genes in mouse zygotes suggest increased transport of transcripts into the cytoplasm from the nucleus, which could also explain observed upregulation of many genes even in the absence of true transcription. On the other hand, upregulated genes in rat zygotes are enriched for the TCA cycle, hinting at a requirement for more energy production after fertilization. Conversely, the TCA cycle genes involved in energy production were downregulated in mice, while the genes involved in acetyl-CoA synthesis from pyruvate were upregulated. These results could be due to differences in the timing of events, whereby the mouse zygote prepares its first mitosis earlier and in a more active manner, while the rat zygote has still yet begun degradation of maternal transcripts. However, our detection of numerous upregulated genes in zygotes relative to oocytes in rats, suggests that MZT might occur in rat zygotes, though at a smaller scale than in mouse zygotes.

Our analysis on intronic transcripts of genes found overall increases in the proportion of intronic reads and number of genes with intronic transcripts in both species after fertilization, indicating an increase of nascent transcription in zygotes. This was particularly true in mice, despite the majority of DEGs found to be downregulated. Previous studies have suggested that the increases in nascent transcription in the zygote is in part due to promiscuous transcription of genes and inefficient splicing during minor zygotic activation in mice^75^, and our results suggest that a similar process might occur in the rat zygotes as well. In addition, coupled with the increased expression activity related to RNA transport, particularly in mice, these changes in nascent and mature transcript proportions may also at least partially result from increased RNA transport from the nucleus to the cytosol.

The observed changes in isoform usage between oocytes and zygotes of both species indicate post-transcriptional regulation of genes that are involved in epigenetic regulation occur in zygotes. It is worth noting that several genes that regulate histone H2A show strong differential isoform usage and differential poly-adenylation in both mice and rats. These results advocate for similar epigenetic regulations of chromatins in both species by differential isoform usage. Though more details remain to be revealed, our results suggest similar post-transcriptional events in zygotes of mice and rats.

Our analysis of differential polyadenylation reveals that transcripts undergo distinct 3’UTR polyadenylations in oocytes and zygotes of both species. We would be remiss to not touch on the issue of differences in gene-body read distribution between oocytes and zygotes and the subsequent impact on alternative polyadenylation analysis. The stronger 3’-end bias of reads in mouse oocytes than in mouse zygotes is unlikely due to technical factors but rather biological differences between mouse oocytes and zygotes, which is not the case in rat oocytes and rat zygotes. Such bias in mouse samples should be negligible on our DAP analysis, since similar 3’UTR preference of the same gene would result in similar bias observed in read coverage (exhibiting similar peaks in the 3’UTR region), while only differences in 3’UTR length preference (caused by differential poly-A cleavage sites) would result in peaks appearing in one group and not in the other. Another issue that has been brought up in previous studies regarding transcriptomic profiles of 1-cell stage zygotes obtained via poly-A enrichment techniques is inflation of expression values due to inherent biases towards longer Poly-A tails in poly-A capture RNA-seq methodologies ^104^. Previous studies have also held conflicting views towards the length of poly-A tails in zygotic transcripts. For example, one study ^105^ reported extensive shortening of poly-A tails in the zygote, while another ^106^ suggested a shift towards longer poly-A tail lengths from MII oocytes to zygotes. Though we were not able to quantify for poly-A tail length in our study, we found an overwhelmingly larger overlap between shortened 3’UTRs and upregulated genes in mouse zygotes, compared to other overlaps between DAP and DEG genes. Yet, we note that there is no correlation between the changes in expression levels and the changes in 3’UTR preference measured by PDUI. Taken together, it is unclear whether the observed upregulation is indirectly caused by changes in poly-adenylation, and it appears that many other genes do not experience significant changes in polyadenylation but are upregulated regardless in mouse zygotes. The case in rats is even more prominent, with most upregulated genes experiencing no significant change in 3’UTR preference.

Although most orthologous genes in mice and rats had similar differential transcriptional patterns in term of DGEs, DNP, DTU and DAP during MZT, others showed different patterns. In summary, these orthogonal analyses on the transcriptomic changes during the oocyte to zygote transition in mice and rats strongly suggest minor zygotic activation is a vibrant process in both species, albeit a delayed and weaker process in rats. Much of the observed results also confirm significant post-transcriptional regulation, most notably in the form of alternative polyadenylation, in both species.

Future studies should involve a multi-omics approach for both species, with strong emphasis on post-transcriptional modifications such as poly-adenylation and epigenetic regulations of expression. Furthermore, the lack of transcript degradation in rat zygotes may imply that timing of rat zygotes and mouse zygotes differ in the progression of MZT. However, there was a much greater number of upregulated DEGs observed in the rat, perhaps owing to transient expressions of the maternal genome upon fertilization. Thus, a more comprehensive analysis could focus on finer-grained snapshots of the zygote in both mice and rats to capture transient expressions of genes, as well as timing of minor zygotic activation.

## Resource availability

### Lead Contact

Further information and requests for resources and reagents should be directed to and will be fulfilled by the lead contact, Zhengchang Su

### Materials availability

No new reagents were generated from this paper.

### Availability of data and materials

- Single-cell RNA-seq data has been deposited at GEO Accession: GSE287856 and is publicly available as of the date of publication.
- All original code has been deposited at https://github.com/bio-info-guy/MiceRatZGA/ and is publicly available as of the date of publication.
- Any additional information required to reanalyze the data reported in this paper is available from the lead contact upon request.

## Acknowledgements

The work was supported by the US National Science Foundation (DBI-1661332) and NIH (R01GM106013). The funding bodies played no role in the design of the study and collection, analysis, and interpretation of data and in writing the manuscript.

