## Supplementary Figures S1 - S4 for "Single-cell RNA-seq Reveals Early Transcriptional Programs of the Maternal to Zygote Transition in Mice and Rats"

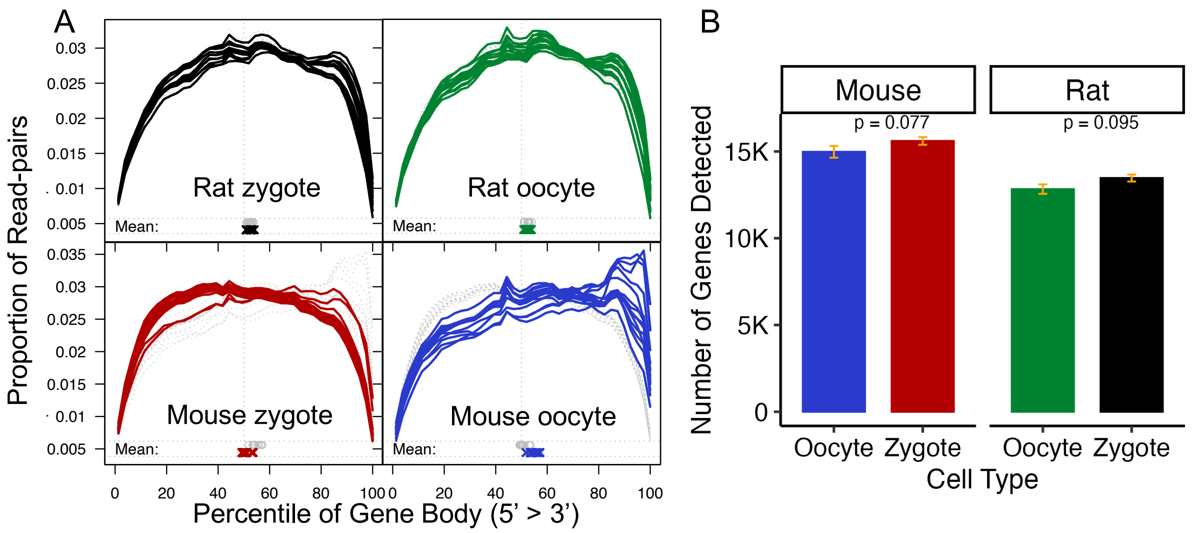


**Supplementary Figure S1** A. Gene-body coverage of all samples separated into cell types. **B.** Number of genes detected by cell type, Wilcoxon rank-sum test p-values.

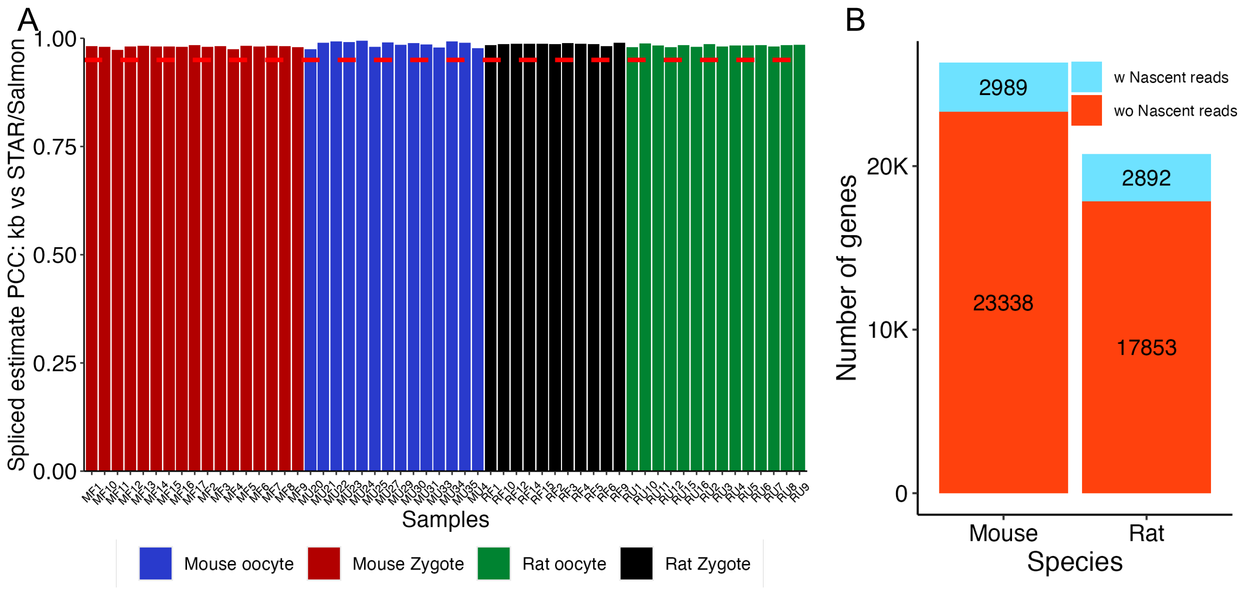


**Supplementary Figure S2 A.** Pearson correlation coefficient (PCC) between kallisto/bustools estimated spliced-reads counts and STAR/salmon estimated read counts for each sample. **B.** Number of genes with nascent read coverage after filtering in each species.

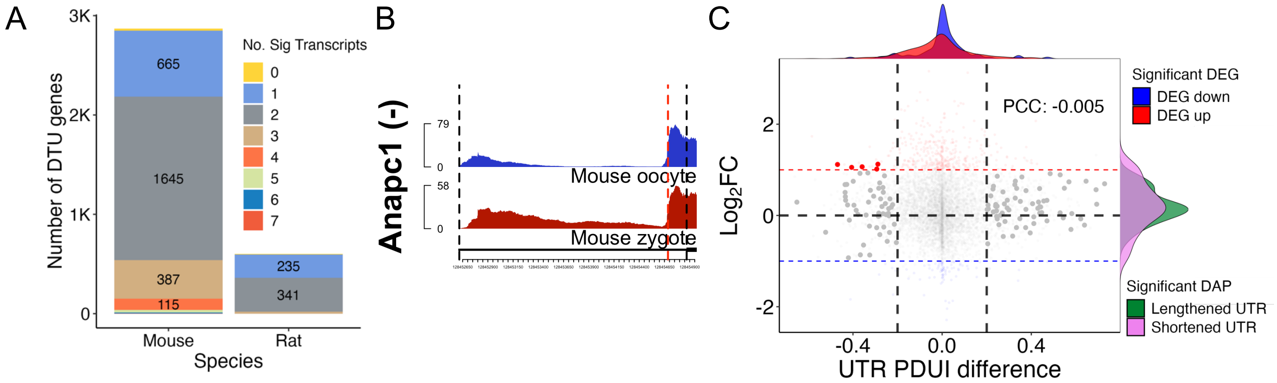


**Supplementary Figure S3 A.** Pearson correlation coefficient (PCC) between kallisto/bustools estimated spliced-reads counts and STAR/salmon estimated read counts for each sample. **B.** 3'UTR coverage of *Anapc1* in mice. **C.** Number of genes with nascent read coverage after filtering in each species.

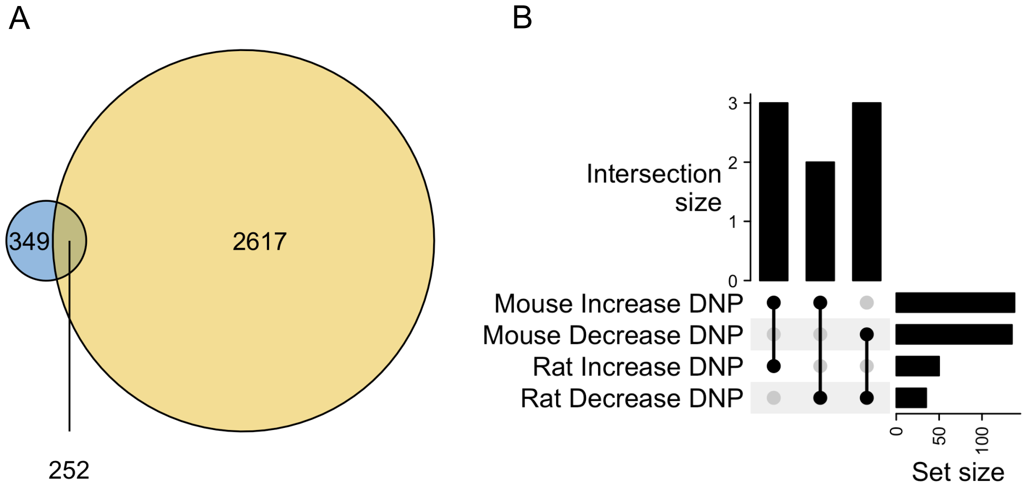


**Supplementary Figure S4 A.** Venn diagram of DTU genes in mice and rats. **B.** Upset plot of DNP genes in mice and rats.
